# Apoptosis at the optic fissure shapes transient retinal myeloid states

**DOI:** 10.64898/2026.10.06.757121

**Authors:** Nathaniel Ghena, Wesley S. Warner, Yesica Landaverde Rodriguez, Joon Schwakopf, Jacqueline M. Roberts, Monica L. Vetter

**Affiliations:** Department of Neurobiology, Spencer Fox Eccles School of Medicine, University of Utah, Salt Lake City, UT; Interdepartmental Program in Neuroscience, University of Utah; Intermountain Doctoral Education to Advance Students (IDEAS) Neuroscience Program, University of Utah

## Abstract

Microglia and CNS-associated macrophages (CAMs) originate from yolk sac erythromyeloid progenitors, yet the process that mediates their colonization of the mammalian eye and the developmental cues that promote their diversification remain unclear. Here, we identify the optic fissure as the primary entry site for myeloid cells into the embryonic mouse retina. This entry is spatially and temporally coupled with a localized wave of cell death. This apoptotic niche contains colonizing myeloid cells in a transient CD206+ state, and apoptotic burden directly correlates with CD206 expression across development. We find that disrupting apoptotic recognition pathways, specifically *Cx3cr1* and *Itgam*, promotes enrichment of CD206+ cells; likewise, increasing cell death via genetic loss of *Atoh7* results in elevated CD206+ population, indicating that impaired clearance and increased death converge on a common outcome within a constrained developmental window. Both CD206+ and CD206- subsets contribute to phagocytosis during optic fissure closure, indicating that CD206 expression does not mark a specialized phagocytic subset. Intriguingly, unlike parenchymal microglia, CD206+ cells are selectively spared from pharmacological CSF1R inhibition, revealing distinct survival requirements within the fissure niche. Together, these findings indicate that localized apoptotic niches encountered during myeloid cell entry influence the initial states and properties of CNS-resident myeloid cells.

## Introduction

The adult central nervous system (CNS) contains two primary myeloid populations: parenchymal microglia, which reside in the neural compartment, and non-parenchymal CNS-associated macrophages (CAMs), which occupy distinct anatomical niches at CNS borders (Prinz et al., 2017; Prinz et al., 2021). Protein expression further distinguishes these populations, with P2RY12 and SALL1 denoting microglia while CD206 (encoded by *Mrc1*) marking CAMs (Dalmau Gasull et al., 2024; Paolicelli et al., 2022). These populations emerge from yolk sac-derived erythromyeloid progenitors and populate the CNS during embryonic development. Originally, these populations were identified as separate lineages fixed early in development (Utz et al., 2020), however, this dogma has recently been challenged suggesting there exists greater plasticity than previously appreciated (Hattori et al., 2023; Scott et al., 2022). Therefore, how these populations become established and colonize tissues, and the extent to which the local environment shapes their states, remain active areas of research.

The developing mammalian eye provides a spatially resolved model to investigate these events. During optic cup formation, the retina undergoes stereotyped morphogenesis as nascent vasculature and boundary structures emerge and resident myeloid cells begin to populate ocular tissues (Ghena, Lopez, et al., 2025; Silverman & Wong, 2018). However, the cellular logic governing how these cells enter the eye and how their state is influenced by local development cues is not well understood. Live imaging in zebrafish demonstrated that myeloid precursors enter the eye through the choroid fissure in association with hyaloid blood vessels (Ranawat & Masai, 2021), yet this route has not been documented in mammals, where classical histological work suggested invasion from vitreous and ciliary margins (Santos et al., 2008). It thus remains unresolved whether a fissure-associated gateway also operates in the mammalian retina, and whether the local environment at that site shapes the state of the cells passing through it.

During mouse eye morphogenesis, closure of the ventral optic fissure occurs between approximately embryonic day 10.5 (E10.5) and E12, accompanied by focal cell death (Braunger et al., 2014; Chan et al., 2020). The apoptotic signals within this transient niche are thus positioned to serve as a local developmental cue. Cell death is pervasive during neurodevelopment and can provide both chemotactic signals and phagocytic targets for incoming myeloid cells (Ashwell, 1991; Ashwell et al., 1989; Bejarano-Escobar et al., 2011; Hardy et al., 2019). We and others have also shown that apoptotic events not only trigger local microglial responses but can shape microglial gene expression programs (Anderson et al., 2022; Anderson et al., 2019; Ghena, Anderson, et al., 2025; Hume et al., 1983; Martineau et al., 2025), raising the possibility that cell death serves as an organizing cue for early retinal immune assembly.

Notably, CD206 is not restricted to mature CAMs. During early brain development, a subset of colonizing myeloid cells express CD206 alongside canonical microglial markers (Hattori et al., 2023; Li et al., 2019; Matcovitch-Natan et al., 2016; Scott et al., 2022), with little evidence that mature parenchymal microglia retain CD206 expression (Masuda et al., 2022; Utz et al., 2020). This raises the possibility that at early stages, rather than marking a committed CAM lineage, CD206 reports a transient, developmentally restricted state adopted by early myeloid cells in response to local cues, although potential cues remain to be defined.

Here we investigated the earliest colonization of the embryonic mouse retina by yolk sac-derived myeloid cells, identifying the optic fissure as their primary anatomical entry portal. This entry is spatially and temporally coordinated with a localized wave of developmental apoptosis. We show that this apoptotic niche is associated with a transient CD206+ state in colonizing myeloid cells that scales with the level of apoptotic cell death and is characterized by relative independence from canonical CSF1R survival signaling. Using genetic receptor knockouts to disrupt apoptotic-cell recognition and removal alongside an *Atoh7-*deficient model with elevated apoptosis, we further show that altering the clearance or microenvironmental burden of cell death recalibrates the proportion of CD206+ myeloid cells without changing overall myeloid abundance. Importantly, this shift in CD206+ proportion is constrained to an early developmental window. Together, our data indicate that developmental cell death is not merely a clearance target but a spatial cue that shapes the recruitment and state of the early ocular immune system.

## Methods

### Animal husbandry and procedures

All animal procedures were conducted in accordance with protocols approved by the University of Utah Institutional Animal Care and Use Committee (IACUC). Mice were housed in an AAALAC accredited animal facility with 12h light/12h dark cycles and ad libitum access to food and water. Both sexes were used for all experiments. To deplete microglia, mice were given PLX3397 (290 mg/kg, AdooQ BioScience, A15520) formulated in standard chow (Research Diets Inc.) from gestational day 8 until gestational day 11, 13, or 14. Timed matings were used to determine embryonic stages, with two-hour timed mating windows and plug detection used to determine gestational ages. Matings set up from 10pm until 12am were collected at 12pm for E9.5, E10.5, and E16.5 timepoints, while matings were set up from 10am until 12pm for collections at 12pm for E9, E10, E11, E12, E13, and E14 timepoints.

### Mouse strains

C57BL/6J (JAX 000664), B6.129P2 (Cg)-*Cx3cr1^tm1Litt^*/J mice (JAX 005582), B6.129X1-*Bax^tm1Sjk^*/J (JAX 002994), B6.129S4-*Itgam^tm1Myd^*/J (JAX 003991), and CBA.Cg-*Csf1r^em1Prdns^*/J (JAX 032783) were all purchased from the Jackson Laboratory. The *Csf1r*^ΔFIRE^ mice (JAX 032783) were backcrossed to C57BL/6J (JAX 000664) to generate heterozygotes, which were subsequently crossed with Csf1r^ΔFIRE^ homozygotes to produce experimental offspring. The B6.129-*Mertk^tm1Grl^*/J and B6.129-*Axl^tm1Grl^*/J strains (Lu et al., 1999) were a kind gift from Dr. Greg Lemke. To generate *Atoh7-*deficient mice, cryopreserved spermatozoa from the B6.129-*Atoh7^tm1Gan^/*Mmucd strain (RRID: MMRRC_042298-UCD) were obtained from the Mutant Mouse Resource and Research Center (MMRRC) at UC Davis. Live colonies were rederived via *in vitro* fertilization performed by the Transgenic & Gene Targeting Mouse Core Facility at the University of Utah.

### Tissue processing

Pregnant females were decapitated following isoflurane asphyxiation, and whole embryos were fixed in 4% PFA for 45 minutes, then washed 3x for 15 minutes in PBS, followed by 12–16-hour consecutive treatments with 15% and 30% sucrose in PBS at 4°C. Embryos were then embedded in OCT compound (Tissue-Tek), stored at -80°C, and sectioned at 50μm thickness.

### Immunostaining

Frozen sections were placed in ice-cold PBS for 10min, post-fixed for 10min in 4% PFA, washed in ice-cold PBS 3 times for 5min each, and then blocked for 1 hr at room temperature (0.2% Triton-X100, 10% BSA, 10% normal donkey serum in 0.01M PBS). Then, they were incubated in primary antibodies at 4°C overnight (0.2% Triton-X100, 5% BSA in 0.01 M PBS), washed 3 times with PBS for 5min each, and incubated in secondary antibodies (5% BSA in 0.01M PBS) with Hoechst 33342 (1:2000, Thermo Fischer Scientific H3570) for 2 hrs at room temperature. Sections were then washed and mounted with Fluoromount-G mounting medium (SouthernBiotech, 0100-01). Antibodies details listed below. Secondary antibodies were produced in donkey against goat, rat, rabbit, guinea pig, and rat IgG, and conjugated to AlexaFluor -405, -488, -555, or -647 (Thermo Fisher Scientific).

### Antibodies Used

Goat polyclonal anti-GFP (1:2000, Abcam ab5450, RRID: AB_304897)

Goat polyclonal anti-CD206 (1:500, R&D Systems AF2535, RRID: AB_2063012)

Rat monoclonal anti-CD68 (1:250, Bio-Rad MCA1957, RRID: AB_322219)

Rat monoclonal anti-CD206 (1:250, Bio-Rad MCA2235, RRID: AB_324622)

Rat monoclonal anti-CD31/PECAM (1:50, BD Pharmingen 550274, RRID: AB_393571)

Rat monoclonal anti-LYVE1 (1:50, Invitrogen 14-0443-82, RRID: AB_1633414)

Rabbit polyclonal anti-IBA1 (1:1000, Wako 019-19741, RRID: AB_839504)

Rabbit monoclonal anti-CC3 (1:500, BD Biosciences 570524, RRID: AB_3661884)

Rabbit monoclonal anti-C1q (1:1500, Abcam ab182451, RRID: AB_2732849)

Rabbit polyclonal anti-PAX2 (1:1000, BioLegend 901001, RRID: AB_2565001)

Rabbit polyclonal anti-P2RY12 (1:400, Millipore Sigma HPA014518, RRID: AB_2669027)

Rabbit polyclonal anti-COLIV (1:100, Millipore Sigma AB756P, RRID: AB_2276457)

Guinea pig monoclonal anti-IBA1 (1:1000, Millipore Sigma ABN1376, RRID: AB_2924932)

### *in situ* hybridization chain reaction (HCR)

Frozen sections were fixed for 15 min in 4% PFA, washed in ice-cold PBS 3 times for 5min each, and then dehydrated in a series of ethanol washes at 50%, 75%, and twice at 100%, then washed in PBS and dried for 5 min. *in situ* hybridization chain reaction was performed as published(Choi et al., 2018) using v3.0 reagents from Molecular Instruments. Samples were treated with Proteinase K for 10mins at 37C, pre-hybridized with 30% probe Hybridization buffer for 30minsat 37C, and then placed in hybridization overnight at 37C with selected probes. Slides were then washed, placed in amplification buffer for 30mins at room temperature, then incubated with hairpins and amplification buffer overnight at room temperature in the dark. Finally, slides were washed, stained with Hoechst, and cover slipped. 10 probes per gene, recognizing all transcript variants for each of *Ms4a7, Apoe, Ctss,* and *C3* were generated by Molecular Instruments.

### Confocal microscopy

Confocal images were acquired on an inverted Nikon A1R Confocal Microscope, with images acquired at 20X objective and 3X digital zoom and 10% overlap multi-point stitching. Z plane stacks were at 0.8µm steps to capture at 0.2µm pixel resolution. Images represent maximal-intensity projections and acquisition settings were consistent across ages and genotypes.

### Image analysis

All counts were blinded and manually performed using Nikon Elements software (Melville, NY). Inner and outer retina boundaries were identified and used for framing the retina parenchyma. Ventral retina was identified through PAX2 staining and by presence of optic fissure, then the retina was demarcated into quarters, with optic fissure in the middle of the ventral quadrant to define ventral versus adjacent retina for quantification. For IBA1+, CD206+, and CC3+ counts, 3-5 sections (50μm thick sections) were averaged for each retina. IBA1+ cell size and CD68 coverage area were measured manually.

### Bulk RNA-seq and microarray analysis

Bulk RNA-seq data from ventral and dorsal retina at E11.5 and E12.5 were generated by Patel et al (Patel et al., 2020) and obtained from ArrayExpress with accession number E-MTAB-9427. Fastq files were trimmed with cutadapt v2.8 and aligned with STAR v2.7.9a in two pass mode to indexed genome GRCm39 using Ensembl release 104. Uniquely aligned fragments were counted with featureCounts v1.6.3. Microarray data of optic fissure, nasal, and temporal retina was generated by Cao et al. (Cao et al., 2018) and obtained from GEO with accession number GSE109501. Affy v1.82.0 was used to process array data with the functions ReadAffy to import signal intensity files and mas5 with normalization off to convert to expression values. Statistical analysis for both datasets was performed with DESeq2 v1.44.0.

Retinal microglia enriched genes were defined by statistical analysis performed in Anderson et al. (Anderson et al., 2019) available in GEO under accession number GSE123757. Retinal microglia genes are those that are significantly enriched in bulk RNA-seq of sorted retinal microglia, when compared to whole retina tissue at both E16.5 and P7. Genes selected for heatmap representation are retinal microglia genes that are significantly enriched in ventral compared to dorsal retina at both E11.5 and E12.5 and are significantly enriched in the optic fissure when compared to both nasal and temporal retina. Gene set enrichment analysis was performed with fgsea v1.36.2 using default parameters. Genes were sorted by fold change in ventral vs. dorsal retina at E11.5 (Patel et al., 2020) or optic fissure vs. temporal retina (Cao et al., 2018) and compared to the mouse MSigDB (msigdbr v26.1.0) GO Biological Process and WikiPathways collections respectively.

## Statistical methods

All image and flow data were analyzed using Prism 10 software (GraphPad, La Jolla, CA). We first tested for normality using four tests: Anderson-Darling, D’Agostino & Pearson, Shapiro-Wilk, and Kolmogorov-Smirnov test. Failure of any one test resulted in non-parametric tests, Mann-Whitney for pair-wise comparisons and Kruskal-Wallis for comparisons of three or more. We tested for heteroscedasticity using an F-test for pairwise comparisons and Brown-Forsythe test for comparisons of three or more. If not significantly different, we either ran a two-tailed T test, Ordinary one-way ANOVA with posthoc Tukey’s multiple comparison test, or Ordinary two-way ANOVA with Sidak’s multiple comparison test. If standard deviations were significantly different between groups, we ran a Welch’s ANOVA with posthoc Dunnett’s T3 multiple comparison’s test. For all data that is presented as the mean, error bars indicate the standard error of the mean, SEM. We used a 95% confidence interval and a *P* value of < .05 was set for rejecting the null hypothesis.

## Data Availability

No new sequencing data were generated in this study. Previously published datasets analyzed here are available from ArrayExpress under accession E-MTAB-9427 (Patel et al., 2020) and from the Gene Expression Omnibus under accessions GSE109501 (Cao et al., 2018) and GSE123757 (Anderson et al., 2019). Raw imaging data are available from the corresponding author upon request.

## Results

### Developmental apoptosis precedes myeloid colonization of the retina

Given the importance of apoptosis as a key developmental cue recognized by myeloid cells, we sought to define the relationship between cell death and colonization of the embryonic mouse retina. To characterize the spatiotemporal dynamics of apoptosis and myeloid entry, we first quantified cleaved caspase-3 positive (CC3+) apoptotic cells from embryonic day 9 (E9) to E14, spanning the period of optic fissure closure (∼E10.5 – E12.5), with tight control for gestational age. We detected CC3+ cells within the retina at E9 and E10, which peaked in number by E11 before declining sharply thereafter (**Figure 1A, B**). At E9, CC3+ cell bodies displayed distinct bimodal distribution, with localization to the ventral retina as well as a sizable group in the dorsal retina likely coinciding with the transient dorsal notch that resolves rapidly during early optic cup morphogenesis (Schook, 1980). Between E10 and E11, we observed apoptotic cell bodies enriched within the margins of the ventral retina at the optic fissure (**Figure 1A, 1B’, 1F**). IBA1+ myeloid cells were first observed in the optic cup at E9.5 (**Figure 1E**), demonstrating that apoptosis precedes myeloid colonization. IBA1+ myeloid cells were highly enriched in the ventral retina during optic fissure closure (E11), then distributed throughout the retina by E13 after closure is complete (**Figure 1A**), with a marked expansion in total IBA1+ cell number from E9.5 through E14 **(Figure 1A, C)**. These data demonstrate that myeloid cells do not colonize the retina uniformly; instead, they utilize the optic fissure as a primary anatomical portal, selectively localizing with regions of high apoptotic cell enrichment.

**Figure 1:**
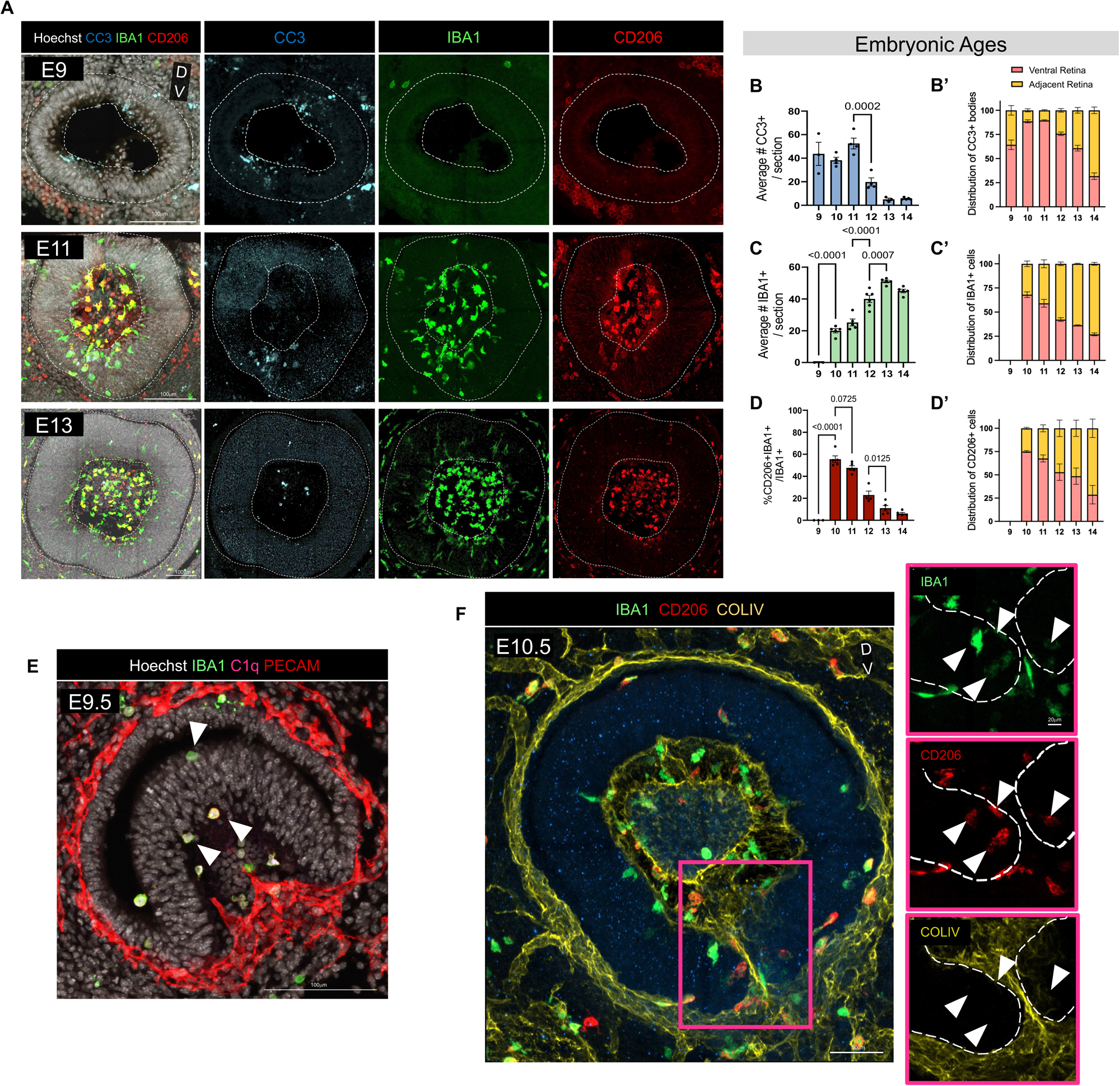
Developmental apoptosis in the ventral optic fissure precedes myeloid cell colonization during retina morphogenesis. (A) Representative retinal sagittal sections of E9, E11, and E13 wildtype mice stained with CC3 (cyan), IBA1 (green) and CD206 (red). Scale bars 100μm. (B) Quantification of average number of CC3+ cell bodies in the retina per section from E9 through E14 (*n =* 3 E9, *n =* 4 E10, *n =* 4 E11, *n =* 4 E12, *n =* 3 E13, *n =* 5 E14). One-way ANOVA F (5, 17) = 25.16 *P*<.0001 and Tukey’s multiple comparisons test. (B’) % of CC3+ cell bodies within ventral retina (pink) versus adjacent retina (yellow) out of total CC3+ cell bodies from E9 until E14 (C) Quantification of average number of IBA1+ cells in the retina per section from E9 through E14 (*n =* 3 E9, *n =* 5 E10, *n =* 5 E11, *n =* 6 E12, *n =* 5 E13, *n =* 5 E14). One-way ANOVA F (5, 23) = 103.0 *P* < .0001 and Tukey’s multiple comparisons. (C’) % of IBA1+ cells within ventral retina (pink) versus adjacent retina (yellow) out of total IBA1+ cells from E9 until E14. (D) Quantification of average number of IBA1+CD206+ cells in the retina per section from E9 through E14 (*n =* 3 E9, *n = 5* E10, *n = 5* E11, *n =* 4 E12, *n = 4* E13, *n =* 5 E14). One-way ANOVA F (5, 21) = 88.17 *P* < .0001 and Tukey’s multiple comparisons test. (D’) % of IBA1+CD206+ cells within ventral retina (pink) versus adjacent retina (yellow) out of total IBA1+CD206+ cells from E9 until E14. (E) Representative image of E9.5 sagittal retina section showing IBA1+C1q+ cells colonizing the retina. IBA1 (green), C1q (pink), PECAM (red). Scale bars 100μm. (F) Representative sagittal section of the ventral retina from E10.5 wildtype mice stained for IBA1 (green), CD206 (red), and COLIV (yellow). Left, merged: the pink box marks the region shown as single channel images. Right, single channel views of the boxed region. Dashed outlines indicate retinal boundaries, white arrowheads indicate IBA1⁺CD206⁺ cells. Scale bar, 50 µm.

### Apoptotic niches serve as a focal hub for CD206+ cells

Recent studies have shown that early colonizing myeloid cells in the embryonic brain can transiently co-express genes associated with both CAMs (*Mrc1*, *Lyve1*) and microglia (*P2ry12*) (Hammond et al., 2019; Hattori et al., 2023; Masuda et al., 2022; Petry et al., 2025; Scott et al., 2022). Given that cell death is spatially restricted to the ventral retina during initial myeloid colonization, we asked whether this apoptotic niche influences myeloid state. At E10 and E11, the majority of myeloid cells (∼60%) within the fissure-associated niche expressed CD206 (encoded by Mrc*1*) (**Figure 1D, 1D’**). These IBA1+CD206+ cells were primarily localized near the inner limiting membrane, often directly apposed to Collagen IV+ basement membranes (**Figure 1F**).

To further characterize these colonizing cells, we examined the expression of canonical microglia marker P2RY12 and CAM marker LYVE1 at E12.5 (**Figure 2A**). We found remarkable phenotypic diversity within the retina, observing subsets of IBA1+ cells that were P2RY12+CD206-LYVE1-, P2RY12-CD206+LYVE1+, and even triple-positive P2RY12+CD206+LYVE1+ within the retina parenchyma (**Figure 2A insets**). This high degree of marker overlap indicates that early-colonizing myeloid cells do not segregate cleanly into CAM and microglia identities, but instead occupy a continuum of intermediate states as they populate the neural retina.

**Figure 2:**
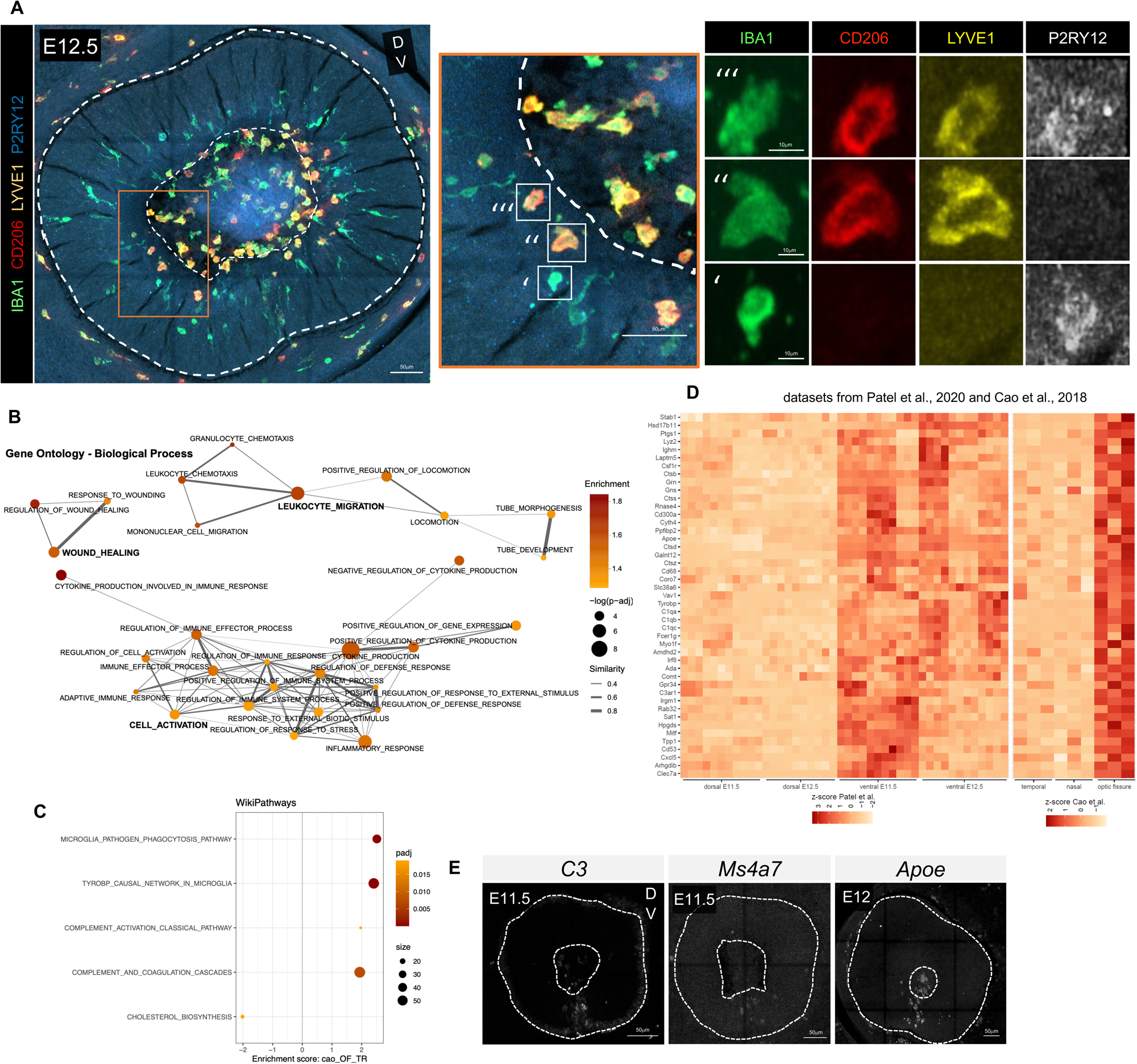
Initial colonizing myeloid cells display both microglia and CAM signatures. (A) Representative sagittal sections of retina at E12.5 stained for IBA1 (green), CD206 (red), LYVE1 (yellow), and P2RY12 (blue). Dashed lines delineate the boundaries of the retinal parenchyma. Left: merged image; Middle: zoomed in orange box; Right, single-channel images for each marker. Inset example representative image of ‘ = IBA1+CD206- LYVE1-P2RY12+ cell; ‘‘ = IBA1+CD206+LYVE1+P2RY12- cell; ‘‘‘ = IBA1+CD206+LYVE1+P2RY12+ cell. Merged image scale bars 50 µm, single channel scale bars 10 µm. (B) The top 30 pathways enriched in the ventral retina at E11.5 (sequencing data from Patel et al., 2020) were plotted as individual points. Point size represents -log(padj), color depicts normalized enrichment score, and line size indicates jaccard similarity of the leading-edge genes for the connected pathways. Similarities of less than 0.25 are not plotted. (C) Top 5 WikiPathways enriched in the ventral optic fissure vs. temporal retina (microarray data from Cao et al, 2018). The size of the gene set considered is represented by point size, color represents significance – (padj), and the x-axis shows the normalized enrichment score. (D) Heatmap of values, scaled by experiment, for myeloid genes significantly enriched in adjacent retina (dorsal, nasal, and temporal) versus ventral retina/optic fissure. (E) Max projection confocal images of *in situ* HCR at E11.5: *C3* and *Ms4a7;* E12: *Apoe*. Retina parenchyma outlined with dashed white line. Scale bars 50μm.

To determine whether the observed spatial differences correspond to distinct functional programs, we analyzed published transcriptomic datasets from microdissected optic fissure and adjacent retina (temporal+nasal+dorsal; Cao et al., 2018; Patel et al., 2020). Gene set enrichment analysis (GSEA) querying gene ontology biological process terms revealed that the ventral retina is significantly enriched for biological processes associated with leukocyte migration, cell activation, and response to wounding (**Figure 2B, Supplementary Table 1**). Consistent with the high demand for debris clearance, GSEA querying WikiPathways showed positive enrichment for the TYROBP causal network in microglia, microglia pathogen phagocytosis, and complement activation pathways in the ventral retina (**Figure 2C**). Furthermore, this region was enriched for genes related to lipid metabolic processes, proteolysis, and lysosomal transport – including cathepsins, matrix metalloproteinases, and ADAMTS family members, aligning with the metabolic requirements of processing apoptotic cargo within the fissure (**Figure 2D**).

We validated these findings using *in situ* hybridization chain reaction (HCR) at E11.5 and E12. We detected robust expression of transcripts encoding for genes related to phagocytosis (*C3* and *Apoe*) and neuro-inflammatory response (*Ms4a7*) that were largely restricted to the ventral retina during fissure closure (**Figure 2E**). Together, these data demonstrate that early colonizing myeloid cells display diverse gene expression profiles, expressing both microglia and CAM markers. Furthermore, CAM gene expression states are enriched in focal apoptotic zones, suggesting that transcriptional identity may be shaped by local developmental cues within the apoptotic niche.

### Both CD206+ and CD206- myeloid cells associate with phagocytosis of apoptotic cells in optic fissure

We next explored the relationship between CD206+ myeloid cells and apoptotic activity. First, we observed a strong negative correlation between CC3+ apoptotic cell bodies and IBA1+ or IBA1+CD206- cell number (**Figure 3A, B**). However, IBA1+CD206+ cells displayed the opposite trend, tracking precisely with apoptotic activity (**Figure 3C**). Thus, the abundance of CD206+ myeloid cells appears to be strongly correlated with the presence of apoptotic cells.

**Figure 3:**
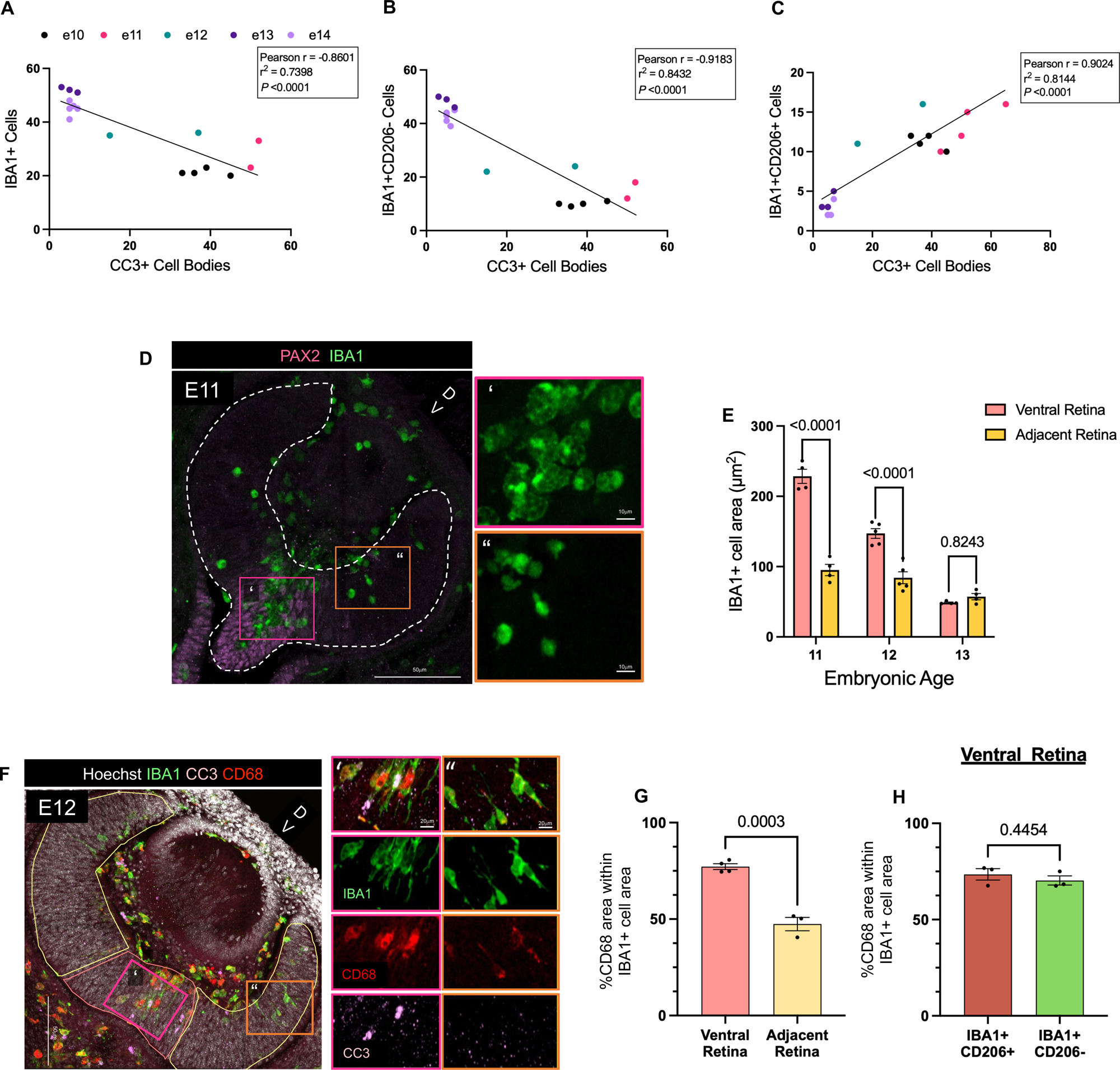
Both CD206+ and CD206- myeloid cells contribute to phagocytosis. (A) Dot plot of average number of CC3+ cell bodies compared to average number of IBA1+ cells per retina of same animal (*n =* 4 E10, *n = 2* E11, *n =* 2 E12, *n =* 3 E13, *n =* 5 E14). Pearson *r* = -0.8601, R squared = 0.7398, *P* < .0001. Average of 3-5 sections/animal. Data points represent mean ± SEM. (B) Dot plot of average number of CC3+ cell bodies compared to average number of IBA1+CD206- cells per retina of same animal (*n =* 4 E10, *n = 2* E11, *n =* 2 E12, *n =* 3 E13, *n =* 5 E14). Pearson *r* = 0.9024, R squared = 0.8144, *P* < .0001. Average of 3-5 sections/animal. Data points represent mean ± SEM. (C) Dot plot of average number of CC3+ cell bodies compared to average number of IBA1+CD206+ cells per retina of same animal (*n =* 4 E10, *n =* 4 E11, *n =* 2 E12, *n =* 3 E13, *n =* 5 E14). Pearson *r* = -0.9183, R squared = 0.8432, *P* < .0001. Average of 3-5 sections/animal. Data points represent mean ± SEM. (D) Representative image of E11 retinal cross-section stained with PAX2 (purple) and IBA1 (green). Pink box ‘ = IBA1 staining in ventral retina, Yellow box ‘‘ = IBA1 staining in adjacent retina. Merged channel image scale bar 50μm, inset scale bars 10μm. (E) Quantification of average number of IBA1+ area (μm^2^) from E11 through E13. (E11: *n =* 4, E12: *n =* 5, E13: *n =* 4). Two-way ANOVA followed by Sidak’s multiple comparisons test. 20-30 IBA1+ cells assessed per ROI/animal, 3 sections averaged / animal / age, 4-5 animals analyzed / age. Bar plot represents mean ± SEM. (F) Representative image of E12 retinal cross-section stained with IBA1 (green), CD68 (red), and CC3 (pink). Pink box ‘ = ventral retina, Yellow box ‘‘ = adjacent retina. Merged image scale bar 50μm, inset scale bars 20μm. (G) Quantification of %CD68 area within IBA1+ cell area (area of CD68+/area of IBA1+) of all IBA1+ cells. Unpaired *t* test. Points represent per animal average. Average of ∼23 IBA1+ cells counted per section, 3 sections averaged / animal, 4 animals analyzed for ventral, 3 animals analyzed for adjacent. Dashed lines represent mean ± SEM of all cells. (H) Quantification of %CD68 area within IBA1+ cell area (area of CD68+/area of IBA1+) for IBA1+CD206+ versus IBA1+CD206- cells. Unpaired *t* test. Points represent per animal average. Average of ∼22 IBA+ cells counted per section, 3 sections averaged / animal, 3 animals analyzed. Bar plot represents mean ± SEM.

Given that CD206+ myeloid cells also occupy the same ventral compartment as CC3+ apoptotic bodies, we asked whether this population was engaged in phagocytic elimination during optic fissure closure. We collected wildtype embryos at E11, E12, and E13 (stages spanning optic fissure closure) and quantified IBA1+ cell morphology since myeloid cells enlarge with phagocytic engagement (**Figure 3D**). We found that IBA1+ cell area was significantly increased in the ventral retina compared to dorsal retina at both E11 and E12. By E13, when fissure closure is complete and CC3+ cell bodies are largely cleared, this morphological difference resolved (**Figure 3E**). To evaluate the phagocytic capacity, we analyzed the expression of the phagolysosomal marker CD68 and observed that CD68 coverage (defined as the ratio of CD68+ area to total IBA1+ area) was significantly enriched in the ventral retina compared to adjacent regions (**Figure 3F, G**), suggesting that myeloid cells within the ventral retina are likely contributing to phagocytic clearance. We then asked whether CD206+ myeloid cells are preferentially phagocytic. Interestingly, when we compared CD68 coverage within the ventral retina, we found no significant difference between CD206+IBA1+ and CD206- IBA1+ populations (**Figure 3H**). These data indicate that while CD206 selectively marks the early myeloid population within the fissure, both myeloid subsets are equally engaged in the clearance of apoptotic debris during optic fissure closure.

### Myeloid cells are required for clearance of apoptotic cells in the fissure

To assess the requirement of myeloid cells in eliminating fissure-closure related cell debris, we pharmacologically ablated CSF1R-dependent myeloid cells, including microglia and some populations of CAMs, using the small-molecule CSF1R inhibitor pexidartinib (PLX3397; **Figure 4A**; Okojie et al., 2023). Continuous administration began at E8, just prior to the onset of yolk sac-derived myeloid circulation (Ginhoux et al., 2010; Kierdorf et al., 2013; Ranawat & Masai, 2021; Schulz et al., 2012; Wu et al., 2018), resulting in >80% depletion of IBA1+ cell by E13 and >95% by E14 (**Figure 4A, B**). Notably, the residual cells were significantly enriched in the ventral apoptotic niche and were predominantly CD206+ (∼75%), suggesting these early myeloid cells are less dependent on CSF1R signaling for survival in the fissure environment (**Figure 4B’, C**). In these depleted embryos, we observed a trend towards accumulation of CC3+ apoptotic bodies by E13, suggesting that the rate of cell death exceeded the clearance capacity of the remaining myeloid population (**Figure 4D**). To corroborate our pharmacological findings with an independent genetic approach, we examined *Csf1r*^ΔFIRE/ΔFIRE^ mice. These harbor a constitutively inactive *Csf1r,* resulting in failure to develop microglia and most CAMs (Rojo et al., 2019). At both E11 and E13, we observed near-complete absence of IBA1+ myeloid cells within the retina and a robust accumulation of CC3+ cell bodies at E13 (**Figure 4E**). This validates our pharmacologic findings that myeloid cells are responsible for the clearance of apoptotic debris during fissure closure. Furthermore, it demonstrates that the initiation of developmental cell death in the optic fissure is cell-autonomous and independent of myeloid cell presence. Interestingly, we see an accumulation not only in the ventral retina, but dorsally as well. This is consistent with impaired clearance of developmental apoptotic material in the absence of microglia, although we cannot exclude a contribution from increased neuronal death secondary to loss of microglial trophic support. Together, these findings indicate that both CD206+ and CD206- cells are essential for maintaining the homeostasis of the apoptotic niche, but do not dictate the emergence of the niche itself.

**Figure 4:**
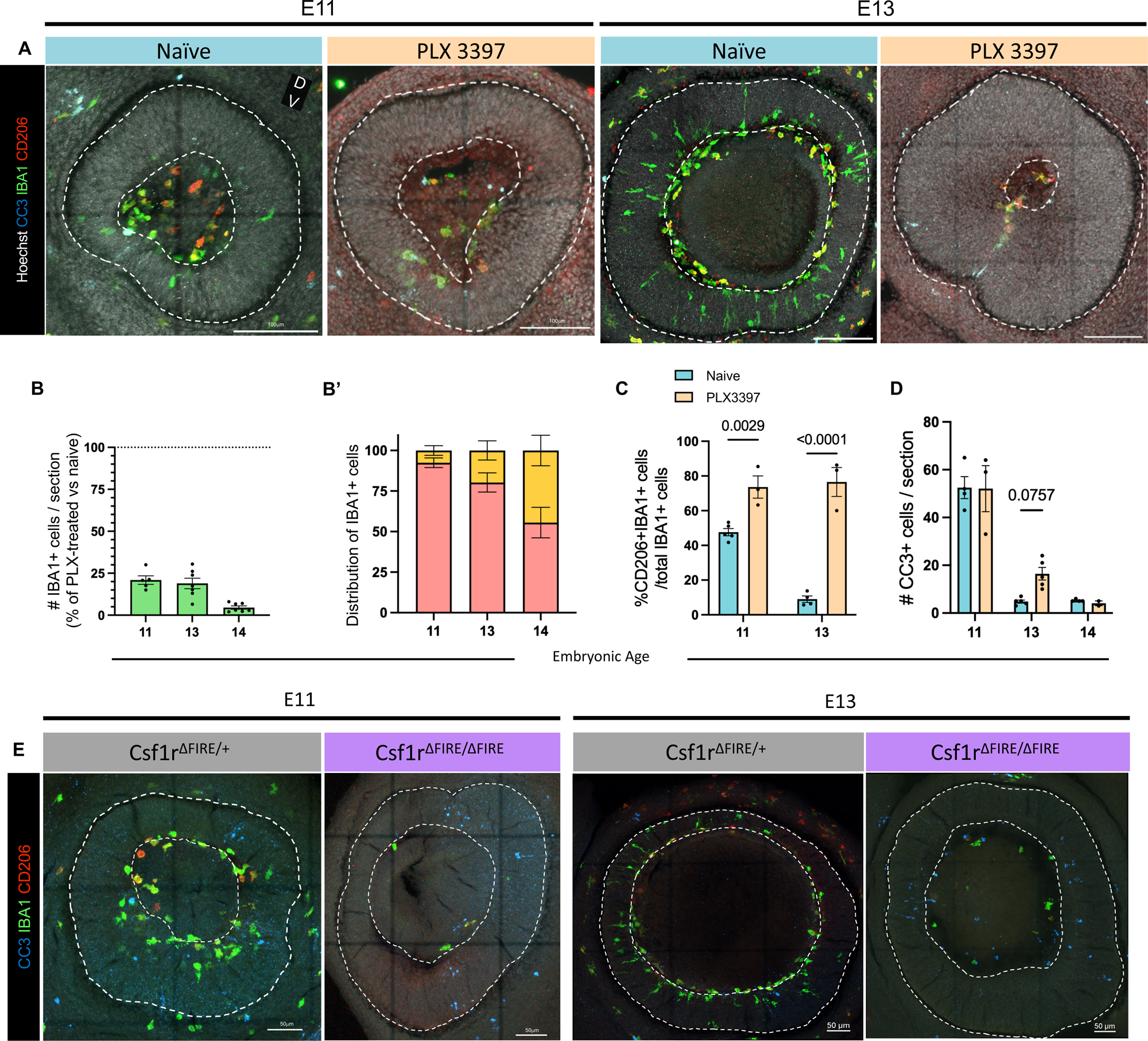
CD206+ cells are less sensitive to acute CSF1R inhibition. (A) Representative images of sagittal retinal sections from naïve and PLX3397-treated wildtype mouse embryos at E11 and E13, stained with CC3 (cyan), IBA1 (green), and CD206 (red). Scale bars 100μm. (B) Quantification of average number of retinal IBA1+ cells remaining after PLX3397 treatment represented as % of controls animals. E11: *n =* 5, E13: *n =* 6, E14: *n =* 7. (B’) % of IBA+ cells within ventral retina (pink) versus adjacent retina (yellow) out of total IBA1+ cells from E11, E13, and E14 naïve and PLX-treated embryos. Average of 3-5 sections/animal/age. Bars and whiskers represent mean ± SEM. (C) Quantification of average proportion of number of IBA1+CD206+ cells / total IBA1+ cells in PLX-treated embryos (orange) versus naïve controls (teal). E11: control *n =* 5, PLX *n =* 3; E13 control: *n =* 4, PLX: *n =* 3. Two-way ANOVA followed by Sidak’s multiple comparisons test. n represents the number of animals, data points are the average of 3-5 sections/animal. Bars and whiskers represent mean ± SEM. (D) Quantification of average number of CC3+ cell bodies / section in retina parenchyma from E11, E13, and E14 PLX-treated embryos (orange) versus naïve controls (teal). E11: control *n =* 4, PLX *n =* 3; E13: control *n =* 5, PLX *n =* 5; E14: control *n =* 4, PLX *n =* 2 PLX. Two-way ANOVA followed by Sidak’s multiple comparisons test. Average of 3-5 sections/animal. Data points represent individual animals, bars and whiskers represent mean ± SEM. (E) Representative images of sagittal retina sections from *Csf1r*^ΔFIRE/ΔFIRE^ embryos and litter mate controls (*Csf1r*^ΔFIRE/+^) at E11 and E13. Merged images depict CC3 (cyan), IBA1 (green), and CD206 (red). Scale bars, 50 μm.

### Apoptotic cell recognition alters CD206+ myeloid cell dynamics

With both CD206+ and CD206- cells involved in phagocytosis but only the CD206+ population tracking with the spatial distribution of cell death, we asked whether localized apoptosis serves as a cellular cue for CD206+ state. To test this, we depleted key genes from molecular pathways known to mediate myeloid recognition and clearance of apoptotic cells: fractalkine receptor *Cx3cr1* (Jung et al., 2000), integrin complement receptor *Itgam* (integrin αM; CD11b/CD18; CR3) (Coxon et al., 1996), and TAM receptors *Mertk* and *Axl* (Lemke, 2017). At E13, we observed no change in total IBA1+ cell density in any of these conditions, suggesting that myeloid cell recruitment to the retina is not altered. *Cx3cr1^-/-^* (CX3CR1 KO) embryos exhibited a significant accumulation of CC3+ cell bodies and a corresponding increase in CD206+ population, while *Itgam^-/-^* (CR3 KO) embryos displayed elevated CD206+ cell counts compared to controls but accumulation of CC3+ cell bodies did not reach significance (**Figure 5A – D**). In contrast, *Mertk^-/-^ (*MERTK KO) and *Axl^-/-^* (AXL KO) embryos showed no change in CC3+ cell body clearance or in the proportion of CD206+ myeloid cells. Our observation that disruption of apoptotic cell recognition or clearance (CX3CR1, CR3) alters CD206+ proportion suggests that the CD206+ state may be induced by dying cells.

**Figure 5:**
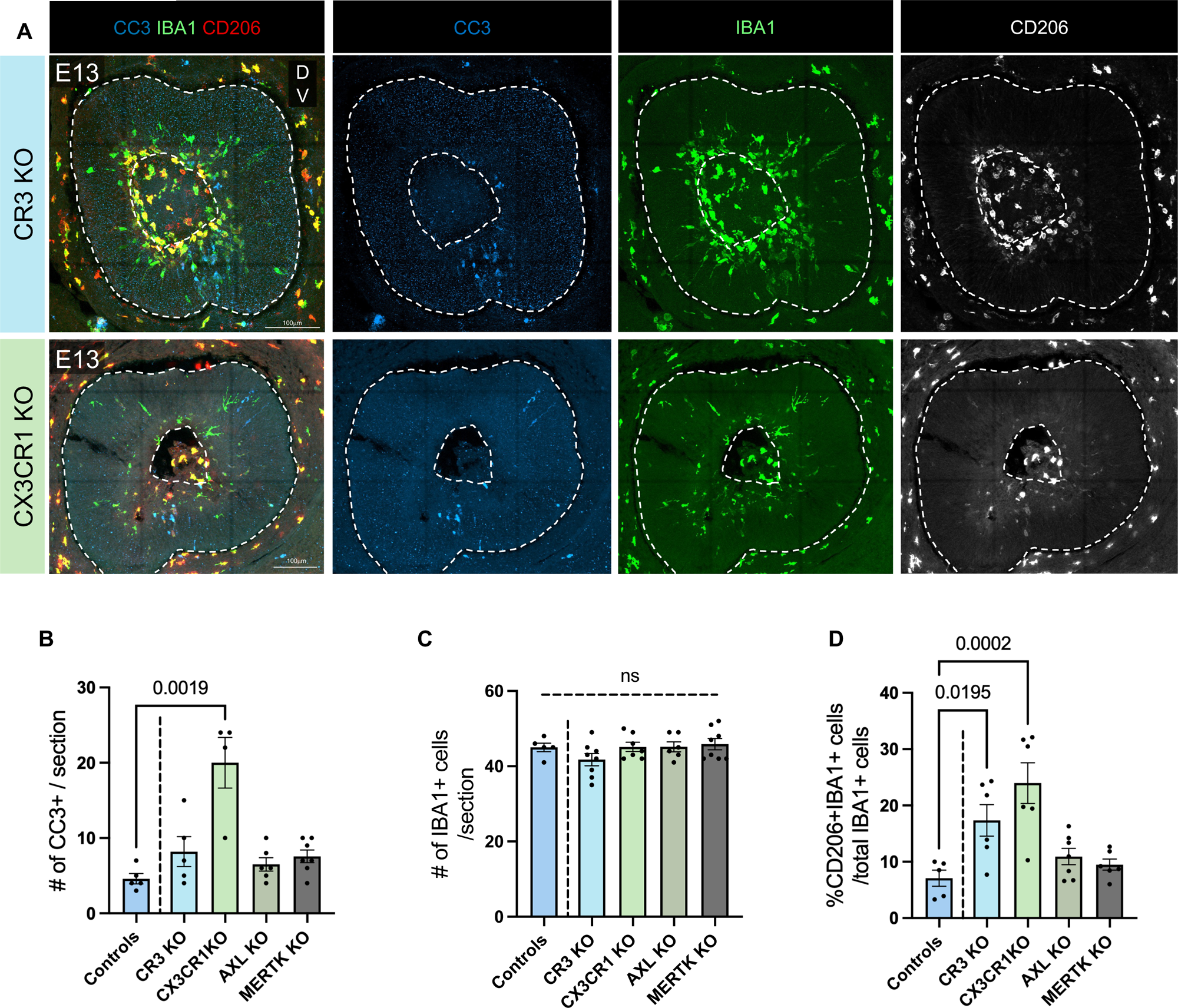
Disruption of cell death recognition pathways CX3CR1 and CR3 increases the proportion of CD206+ microglia. (A) Representative image of sagittal E13 retina from *Cr3* and *Cx3cr1* knockout mice stained with CC3 (cyan), Iba1 (green), and CD206 (red or white). Scale bars 100μm. (B) Quantification of average number of CC3+ cell bodies / section (Control: *n =* 8, *Itgam* KO: *n =* 4, *Cx3cr1* KO: *n =* 3, *Axl* KO: *n =* 6, *Mertk* KO: *n =* 7). Kruskal-Wallis test with Dunn’s multiple comparisons tests. Average of 3-5 sections/genotype. Bars and whiskers represent mean ± SEM. (C) Quantification of average number of IBA1+ cells / section (Control *n =* 5, *Itgam* KO *n =* 8, *Cx3cr1* KO *n =* 7, *Axl* KO *n =* 6, *Mertk* KO *n =* 8). One-way ANOVA F (4, 29) = 1.450 *P =* .2428 and Tukey’s multiple comparisons. Average of 3-5 sections/genotype. Bars and whiskers represent mean ± SEM. (D) Quantification of average proportion of IBA1+CD206+ cells / total IBA1+ cells (Control: *n =* 5, *Itgam* KO: *n =* 6, *Cx3cr1* KO: *n =* 6, *Axl* KO: *n =* 7, *Mertk* KO: *n =* 6). One-way ANOVA F (4, 25) = 8.618 *P =* .0002 and Tukey’s multiple comparisons test. Average of 3-5 sections/genotype. Bar and whiskers represent mean ± SEM.

To ask whether increasing apoptotic load could similarly drive this phenotype, we used *Atoh7* (*Math5*) knockout embryos. *Atoh7* is a bHLH transcription factor required for normal retinal ganglion cell (RGC) fate specification (Brown et al., 2001; Wang et al., 2001). In its absence, RGC-like cells are generated but the majority undergo CC3+ programmed cell death, with >95% of cells that would have become mature RGCs eliminated shortly after specification (Brodie-Kommit et al., 2021). This provides a genetic means to elevate developmental apoptosis without directly manipulating myeloid cells. At E13, *Atoh7^-/-^* embryos exhibited a robust increase in CC3+ apoptotic bodies (**Figure 6A, B)**. Consistent with our hypothesis, we observed this increased apoptotic load coincided with increased CD206+ myeloid cells, but no increase in total density of IBA1+ myeloid cells (**Figure 6A – D**). To determine whether this also occurs at later developmental stages, we examined *Atoh7^-/-^* retinas at E16.5. Despite a continued, massive presence of CC3+ cell death and a marked increase in IBA1+ cells at this later stage, we observed no significant change in the proportion of cells expressing CD206 compared to controls (**Figure 6A – D**). This indicates that at early embryonic stages, myeloid cell interaction with apoptotic cells shifts the proportion that are CD206+ without altering total myeloid cell density, but that this is a transient event limited to the initial stages of retinal colonization. Collectively, these results suggest that CD206 expression in early colonizing retinal myeloid cells reflects a phenotypic state induced by the early developmental environment.

**Figure 6:**
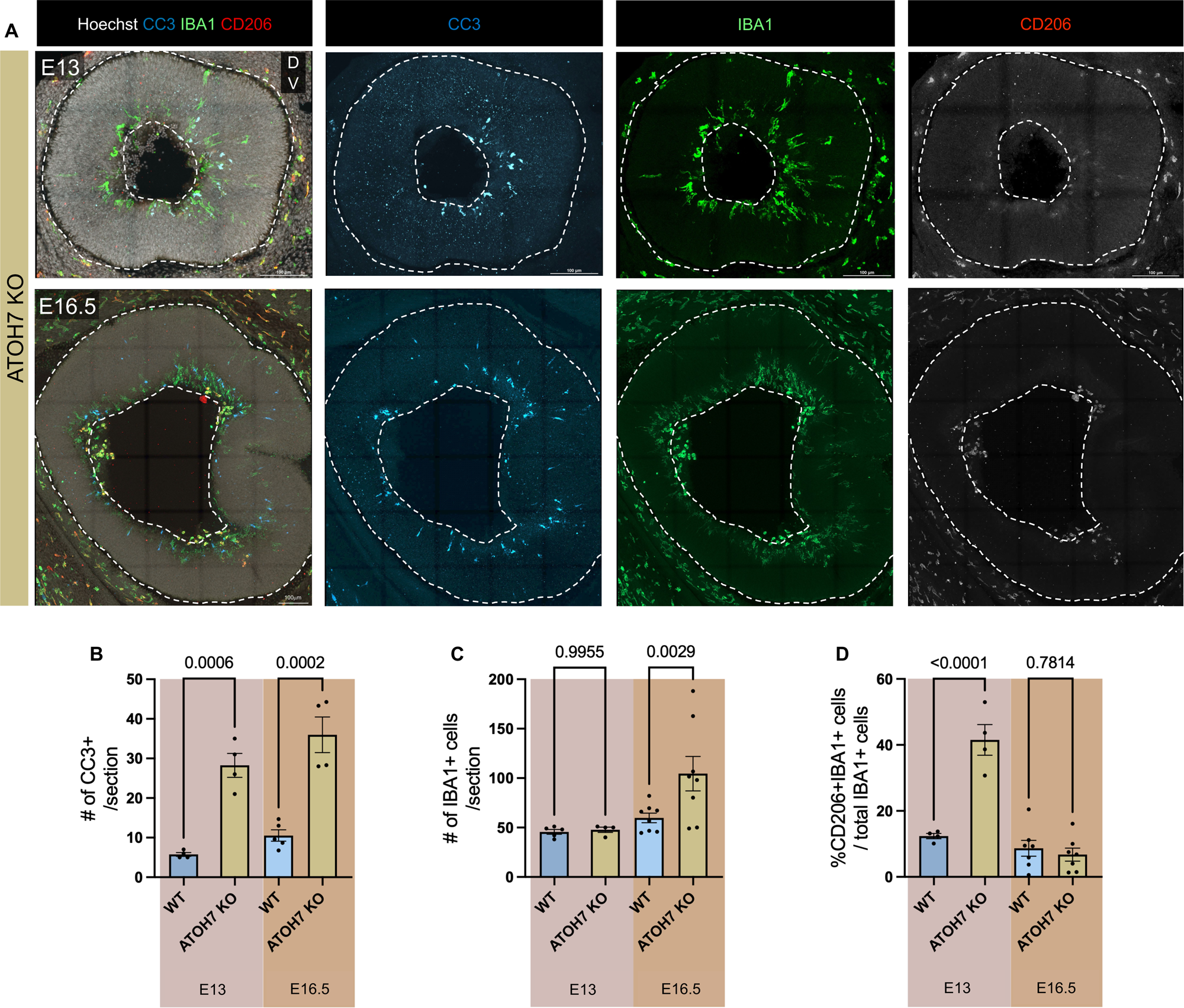
Excess retinal ganglion cell death shifts proportion of CD206+ microglia in a temporally constrained manner. (A) Representative images of sagittal retinal sections of *Atoh7* knockout mice at E13 (TOP) and E16.5 (BOTTOM) stained with CC3 (cyan), IBA1 (green), CD206 (red or white) and Hoechst (white). Dashed line delineates the retina boundary. Scale bars 100μm. (B) Quantification of CC3+ cell bodies / section from control and *Atoh7* KO animals at E13 and E16.5. (E13: control *n =* 4, *Atoh7* KO *n =* 4; E16.5: control *n = 5, Atoh7* KO *n = 4*). Two-way ANOVA followed by Sidak’s multiple comparisons test. Average of 3-5 sections/animal. Bars and whiskers represent mean ± SEM. (C) Quantification of average number of IBA1+ cell / section from E13 and E16.5 control and *Atoh7* KO animals. (E13: control *n =* 5, *Atoh7* KO *n =* 4; E16.5: control *n = 8, Atoh7* KO *n = 8*). Two-way ANOVA followed by Sidak’s multiple comparisons test. Points represent average of 3-5 sections/animal. Bars and whiskers represent mean ± SEM. (D) Quantification of average proportion of IBA1+CD206+ cells / total IBA1+ cells from E13 and E16.5 controls and *Atoh7* KO animals. (E13: control *n =* 4, *Atoh7* KO *n =* 4; E16.5: control *n = 7, Atoh7* KO *n = 7*). Two-way ANOVA followed by Sidak’s multiple comparisons test. Average of 3-5 sections/condition. Bars and whiskers represent mean ± SEM.

Finally, we sought to test whether suppressing apoptosis would conversely reduce CD206+ cell accumulation using *Bax^-/-^* (BAX KO) embryos. While BAX is a requisite pro-apoptotic factor for postnatal neuronal survival, we did not detect a significant difference in CC3+ burden, IBA1+ density, or CD206+ cells at E11 in the retina of BAX KO embryos (**Supplementary Figure 1A – C**). This suggests that the early developmental apoptosis associated with fissure closure may be regulated through BAX-independent pathways (Noh et al., 2016), further emphasizing the specialized nature of this transient niche. Collectively, our findings reveal that the CD206+ myeloid state in the embryonic retinal parenchyma is transiently induced by localized interactions with apoptotic debris.

## Discussion

In this study, we identify the optic fissure as the primary entry portal for myeloid cells into the embryonic mouse retina and show that developmental apoptosis provides a critical spatial cue that orchestrates their recruitment and transient specialization during retina colonization. We find that cell death precedes the onset of colonization and spatially coordinates the entry of yolk sac-derived progenitors through the optic fissure. These early colonizing cells transiently display CD206+ state that scales with apoptotic burden and is characterized by distinct trophic requirements, but are not specialized in their phagocytic activity. Together, these findings reveal that focal apoptosis defines a transient developmental niche that links nervous system morphogenesis with the functional assembly of the innate immune system.

### Optic Fissure as a Portal for Myeloid Cell Colonization of the Eye

Prior studies of mouse retina development reported IBA1+ cells present in the retina by E11.5, but we found evidence for myeloid cells in the retina by E9.5. This earlier timeline aligns with the well-documented initial waves of microglial colonization in other CNS regions which often occur concomitantly with early embryonic vascularization (Arnold & Betsholtz, 2013; Ginhoux et al., 2013). Notably, our data establish that the ventral optic fissure serves as a major anatomical gateway for myeloid cells to colonize the developing mammalian eye. This is consistent with *in vivo* imaging studies in zebrafish and developmental tracing in chicks, which have demonstrated that primitive macrophages utilize the choroid/optic fissure to gain access to the optic cup (Cuadros et al., 1993; Ranawat & Masai, 2021). While alternative secondary entry routes have been noted, including the ciliary margins that supply a distinct contribution of myeloid cells to the peripheral retina (Santos et al., 2008), our findings underscore the ventral fissure as the predominant entry axis during initial ocular seeding.

### Developmental Apoptosis and Myeloid Cell Colonization

The observation that myeloid recruitment coincides with localized apoptosis raises important questions about how dying cells signal to migrating cells. IL-34/CSF1R signaling is well established as a trophic pathway supporting microglial survival and colonization (Easley-Neal et al., 2019; Elmore et al., 2014; Wu et al., 2018), yet our data and others indicate additional cues act in trans to direct regional recruitment. In zebrafish, colonization of the optic tectum is guided by apoptotic neurons releasing lysophosphatidylcholine, which signals through receptor *gpr132b* on myeloid cells (Xu et al., 2016). However, the requirements for cell death appears context-dependent, as apoptosis is dispensable for colonization of the zebrafish retina (Ranawat & Masai, 2021), whereas myeloid entry into the rat retina and forebrain closely tracks with apoptotic events (Ashwell, 1991; Ashwell et al., 1989).

The coupling of myeloid recruitment to localized apoptotic niches may reflect a general principle of CNS development. Early CD206+ myeloid populations have been observed following similar temporal patterns in the embryonic brain and spinal cord (Hattori et al., 2023; Lawrence et al., 2024; Matcovitch-Natan et al., 2016; Scott et al., 2022; Wlodarczyk et al., 2017), although the cues involved were not defined. We propose that the CD206+ state in the embryonic parenchyma represents a transient niche-adapted phenotype that can be induced when myeloid cells encounter a high-demand apoptotic environment during organogenesis. This is unlikely to be the only context in which CD206 is expressed: CNS-associated macrophages maintain CD206 constitutively at CNS interfaces in the absence of substantial cell death, and CD206 itself binds collagen and other extracellular matrix components (Martinez-Pomares, 2012), raising the possibility that the basement membrane contact contributes to its expression or retention. Our data suggest that apoptotic burden scales the proportion of CD206+ cells within the parenchyma, rather than that apoptosis is the sole determinant of CD206 expression. By linking apoptotic patterning to the spatial and molecular assembly of these populations, our findings reframe retinal colonization as a dynamic environmentally guided process in which the first resident immune cells are shaped by the morphogenetic events they are recruited to resolve.

### Apoptotic Cell Recognition Drives the Prevalence of the CD206+ State

The requirement for apoptotic cell recognition to shift the proportion of CD206+ myeloid cells further suggests a central role for the apoptotic niche. Loss of either CX3CR1 or CR3 increased the proportion of CD206+ myeloid cells without altering total myeloid density, indicating a shift in cell state rather than in recruitment. Although disrupting recognition might be expected to blunt an apoptosis-driven signal, the two knockouts act through distinct routes: we found that CX3CR1 loss elevated the unresolved apoptotic burden, whereas CR3 loss shifted the state with little change in burden, indicating that the CD206+ state is in response to both the amount of apoptotic material and how it is sensed. In contrast, loss of MERTK or AXL had no detectable effect, suggesting functional redundancy among TAM receptors (Fourgeaud et al., 2016; Lemke, 2017) or alternate pathways being deployed. This is distinct from postnatal retina, where MERTK, but not AXL, is required for clearance of apoptotic retinal ganglion cell death (Anderson et al., 2022; Lopez et al., 2025). Notably, we found that CD206+ and CD206- cells similarly engulf apoptotic material (Figure 3H), indicating that CD206 does not mark a phagocytically specialized subset, despite previous findings implicating CD206 in phagocytosis (Azad et al., 2014). Together with the expansion of CD206+ cells under genetically increased apoptotic burden in *Atoh7* mutants, these results argue that the CD206+ state is a graded response to the apoptotic environment, or a rapidly changing transient state rather than a division of labor among functionally distinct myeloid populations.

### CSF1R-Independent Survival

One of the more striking findings of our study is the selective survival of CD206+ cells during pharmacological CSF1R inhibition. While most of the early retinal myeloid population was eliminated by PLX3397, the cells residing within the apoptotic niche were significantly spared. This suggests that the fissure environment provides alternative trophic support that bypasses the canonical CSF1R survival pathway. Because both CD206+ and CD206- cells engulf apoptotic material (Figure 3H), selective survival is unlikely to reflect a cell-intrinsic phagocytic advantage. We therefore favor a model in which the fissure niche itself confers trophic support, whether through locally elevated IL-34, the abundance of apoptotic-derived metabolites available to any cell in that location, or other fissure-derived cytokines. Prior work supports the existence of CSF1R-independent survival of microglia in the postnatal retina, as well as developing white matter tracts of the brain during periods of high developmental apoptosis (Anderson et al., 2019; Ghena, Anderson, et al., 2025; Wlodarczyk et al., 2017). Although microglia maintain *Csf1r* expression, we showed that the majority of postnatal retinal microglia survive independently of CSF1R while neuronal developmental apoptosis is high (Anderson et al., 2022). Further work is required to better understand why there is selective independence from CSF1/IL-34 signaling when microglia interact with developmental apoptotic death.

### Niche-Driven Myeloid Plasticity

Historically, the expression of CD206 (*Mrc1*) in the CNS was thought to exclusively define CNS-associated macrophages (CAMs) as a distinct lineage from P2RY12+ microglia (Utz et al., 2020). Our findings, in conjunction with others (Hattori et al., 2023; Masuda et al., 2022; Petry et al., 2025; Scott et al., 2022)], challenge this rigid ontogenetic model, demonstrating instead that the CD206+ state is a plastic, environment-dependent state for early myeloid cells. We observe that CD206 and LYVE1 expression is concentrated among fissure-associated cells and diminishes across the parenchyma, inversely mirroring the distribution of P2RY12, with intermediate cells transiently co-expressing all three markers. Because these data are cross-sectional, we cannot formally exclude that CD206⁺ and P2RY12⁺ cells arise as independently recruited populations; however, the continuous marker gradient and the co-expressing intermediates we observe favor a model in which colonizing cells progressively downregulate the CAM-associated program as they mature within the parenchyma. This transition likely reflects a competitive hierarchy of environmental cues. While the high apoptotic load of the fissure induces and sustains the CD206+ state, the maturing retinal parenchyma provides dominant TGFβ-dependent signals that progressively override it. Recent evidence establishes that TGFβ signaling, mediated by SMAD4, is the primary driver of the core microglial program (Brioschi et al., 2023; von Ehr et al., 2020). This pathway directly regulates the expression of the transcriptional repressor *Sall1* (Brioschi et al., 2023; Fixsen et al., 2023), which is essential for silencing non-microglial myeloid genes. Notably, loss of *Sall1* results in the aberrant persistence of CD206 expression into postnatal ages (Scott et al., 2022). Our observation in *Atoh7* mutants at E16.5, where cells fail to maintain CD206 expression despite a massive continued apoptotic burden, supports the existence of a ‘temporal gate’ in myeloid plasticity. We propose that once the TGFβ/SALL1 maturation program is engaged, the cell becomes refractory to apoptotic cues, enforcing a terminal microglial identity.

Overall, our findings establish that developmental apoptosis within a focal niche in the optic fissure promotes a transient CD206+ state of myeloid cells enriched in this region. The dynamic scaling of this state with apoptotic burden highlights the plasticity of embryonic myeloid cells and reveals how neural morphogenesis coordinates assembly of the resident innate immune system.

## Acknowledgements

This research was supported by the National Eye Institute and the National Institute of Neurological Disorders and Stroke of the National Institutes of Health, under awards R01EY030307 and R21 EY034753 (MLV), R25NS130964 (YR), and T32EY024234, T32NS115664, F31EY035163 (NG) and by the National Science Foundation Graduate Research Fellowship Program (WW). We thank the University of Utah BPRB Animal Facility.

## Supplemental Figure Legends

**Supplementary Figure 1: Related to Figure 6.**
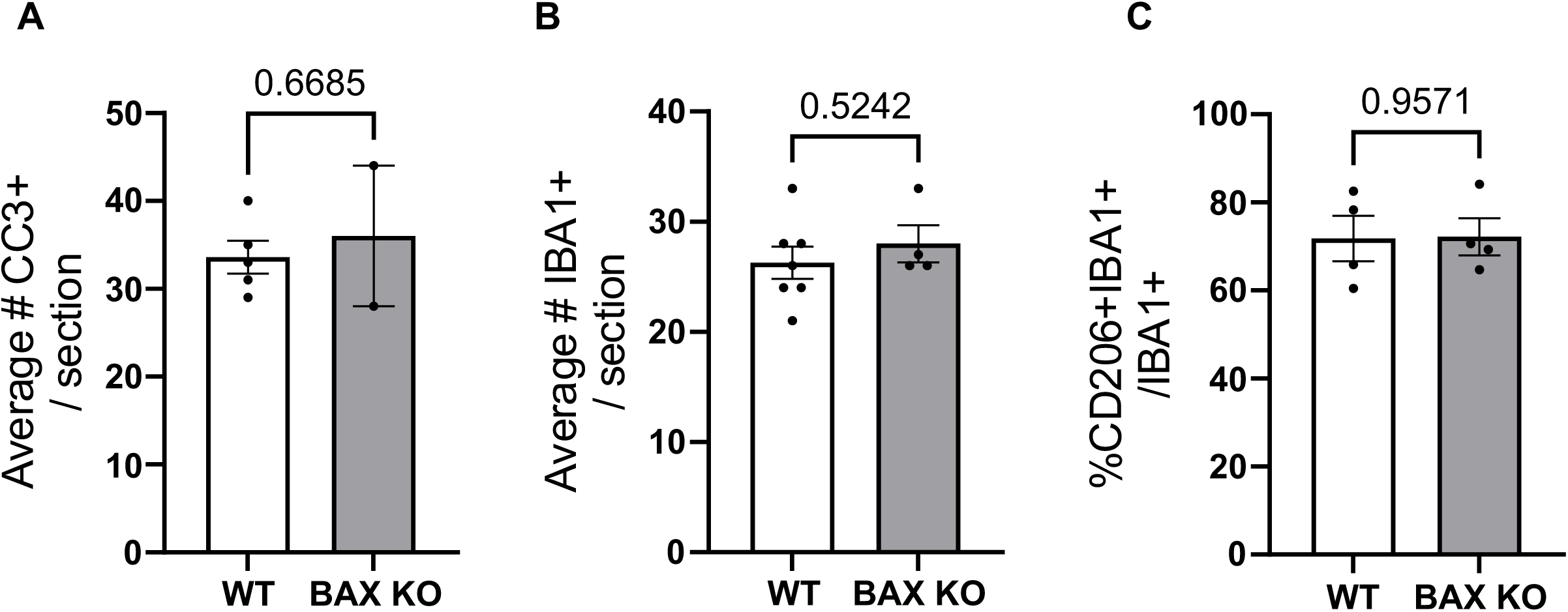
(A) Quantification of average number of CC3+ cell bodies / section from E11 controls and *Bax* KO retinas. Control: *n =* 5, *Bax* KO: *n =* 2. Unpaired *t* test. Average of 3-5 sections/animal. Bars and whiskers represent mean ± SEM (B) Quantification of average number of IBA1+ cells / section from E11 controls and *Bax* KO retinas. Control: *n =* 7, *Bax* KO: *n =* 4. Unpaired *t* test. Average of 3-5 sections/animal. Bars and whiskers represent mean ± SEM (C) Quantification of average number of IBA1+CD206+ cells / total IBA1+ cells / section from E11 controls and *Bax* KO retinas. Control: *n =* 4, *Bax* KO: *n =* 4. Unpaired *t* test. Average of 3-5 sections/animal. Bars and whiskers represent mean ± SEM

